# Pre-existing Immunity to Influenza Aids Ferrets in Developing Stronger and Broader Vaccine-induced Antibody Responses

**DOI:** 10.1101/2024.01.31.578203

**Authors:** Yang Ge, Yao Lu, James Allen, Tal Einav, Dennis Iziogo Nkaleke, Fengwei Bai, Andreas Handel, Ted Ross, Ye Shen

## Abstract

Influenza seasons occur annually, building immune history for individuals, but the influence of this history on subsequent influenza vaccine protection remains unclear. We extracted data from an animal trial to study its potential impact. The trial involved 80 ferrets, each receiving either one type of infection or a placebo before vaccination. We quantified the vaccine protection by evaluating hemagglutination inhibition (HAI) antibody titer responses. We tested whether hosts with different infection histories exhibited similar level of responses when receiving the same vaccine for all homologous and heterologous outcomes. We observed that different pre-existing immunities were generally beneficial to vaccine induced responses, but varied in magnitude. Without pre-immunity, post-vaccination HAI titers after the 1st dose of the vaccine were less likely to be above 1:40, and a booster shot was needed. Our study suggests that pre-existing immunity may strengthen and extend the homologous and heterologous vaccine protection.

## Introduction

Influenza causes millions of infections and hundreds of thousands of deaths each year [1, 2], creating natural immunity in survivors. This immunity, along with the one induced by vaccination, forms a complex history of pre-existing immunity in the human population that is different for every individual [3].

The role of pre-existing immunity is crucial in determining the strength and breadth of future vaccine protection; however, the direction and magnitude of this impact remain unclear. Regarding the accumulating immune history for each individual, some researchers claim that this could be “original antigenic sin,” which suggests that the virus causing the very first influenza infection may establish a dominant memory in the human immune system, diverting the immune response and potentially distracting the protection from against infections caused by other influenza viruses [4, 5, 6]. On the other hand, others consider it as antigenic seniority or antigen imprinting, which offers cross-protection when a new strain emerges [7, 8].

The complex infection and vaccination history among the human population has posed extreme difficulties in exploring the impact of pre-existing immunity [9]. Therefore, animal models have gained popularity as an alternative approach to study this phenomenon [10, 11, 12]. Thus, studying vaccine-induced protection among ferrets with pre-existing immunity due to infection can provide evidence about the interaction between immune history and vaccination that contributes toward future protection [13].

In this study, we extracted data from one ferret trial that implemented four different influenza virus challenges before vaccination, then administered five different vaccines [14]. Uniquely, to better explore the impact on homologous and heterologous protection, 25 historical and 9 co-circulating H3N2 viral strains were used to evaluate the antibody responses generated in each pre-immune setting. We found that pre-existing immunity benefited the induced vaccine mediated protection.

## Methods

We aimed to investigate the heterogeneity in the strength and breadth of vaccine-induced immunological responses related to pre-vaccination infection histories. To achieve this objective, we conducted a secondary data analysis based on a previous animal trial completed by Allen et al. [14].

### Study Design

The animal trial involved 80 ferrets. Before the later vaccination, each was infected with either one type of influenza virus or a placebo (mock). Subsequently, they were administered one of four influenza vaccines or a placebo (mock). With four infection options and five vaccine options (4*5=20), each scenario had four ferrets (4*20=80). The trial aimed to mimic the immune history of humans in different birth cohort by providing ferrets with preexisting immunity before vaccination. The influenza A(H3N2) viral infection strains given before vaccinations were (1) A/Panama/2007/1999 (Pan/99), (2) A/Sichuan/2/1987 (Sich/87), (3) A/Port Chalmers/1/1973 (PC/73), and (4) Mock [14]. Two new vaccines, TJ-2 and TJ-5, were tested in the trial. TJ-2 was designed using H3N2 wild-type hemagglutinin (HA) sequences from the years 2002-2005, and TJ-5 was created using H3N2 sequences from the years 2008-2012 [15]. The next-generation methodology called COBRA (computationally optimized broadly reactive antigen) was implemented to build these two vaccines [15]. Thus, the vaccines used in this trial were (1) A/Wisconsin/67/2005 (Wisc/05), (2) TJ-2, (3) A/Texas/50/2012 (Tx/12), (4) TJ-5 and (5) Mock [14].

The influenza virus was inoculated into each animal on Day 0. The first vaccine/placebo was administered on Day 84, and the second vaccine/placebo, identical to the first, was given on Day 168. Blood samples were collected on Day 14 (14 days after virus exposure), and on Days 98 and 182 (14 days after each vaccination). For each sample, hemagglutination inhibition (HAI) assay responses were tested against 47 different H3N2 viruses (25 historical and 22 co-circulating strains). The HAI titer data were produced by HAI assays following standard protocols [14].As some strains were only tested after one of the two vaccinations, we had to exclude them, resulting in our final dataset incorporating HAI titers for 25 historical and 9 co-circulating strains (detailed strain names provided in the appendix).

### Data Processing

To study the strength (homologous response against the vaccine strain) and breadth (heterologous response against other strains) of the vaccine-induced immunity, we utilized log2 transformation for HAI titers. Because COBRA TJ-2 and TJ-5 are made using computationally optimized broadly reactive antigens [16], they did not have a clear definition of homologous response, as no single strain was the same as the vaccine. Similarly, the mock also did not have a homologous response. Therefore, only the Wisc/05 and Tx/12 vaccines had homologous HAI data. We measured the HAI heterologous responses with the geometric mean across all strains, separated by historical and co-circulating strains.

We quantified the impact of the vaccine on HAI titers using two common outcomes: (1) Titer increase [17], (2) Post-vaccination titer [18]. The titer increase was defined as the difference between post-vaccination and pre-vaccination titers in the log2 scale.The titer increase after the 1st vaccine was defined as (log_2_ HAI_*Day*98_ − log_2_ HAI_*Day*14_). The 2nd titer increase was defined as (log_2_ HAI_*Day*182_ − log_2_ HAI_*Day*98_).

### Statistical models

Our aim is to answer the question of whether hosts with different infection histories will exhibit a similar level of antibody responses when receiving the same vaccine. Therefore, we compared the HAI titer values after one vaccine across all different pre-immunity groups. To assess vaccines, we also compared the antibody levels of different vaccine groups with the same pre-immunity. These values were compared using the Wilcoxon or Kruskal-Wallis tests for all homologous and heterologous outcomes [19]. The Kruskal-Wallis test is a non-parametric statistical test used to determine if there are significant differences between three or more independent groups or treatments. It is often employed when the assumptions of a one-way ANOVA (Analysis of Variance) test, a parametric test, are not met. Finally, we used the Holm-Bonferroni method for pairwise comparisons [20]. All models and analyses were implemented in R 4.3.2 [21].

## Results

In this study, 80 ferrets were assigned to 20 trial groups. Each group received different infections with influenza viruses and vaccines. Their serum samples were collected and used to assess HAI titers against 25 historical (Figure 2) and 9 co-circulating influenza strains (Figure 3), respectively.

**Figure 1:**
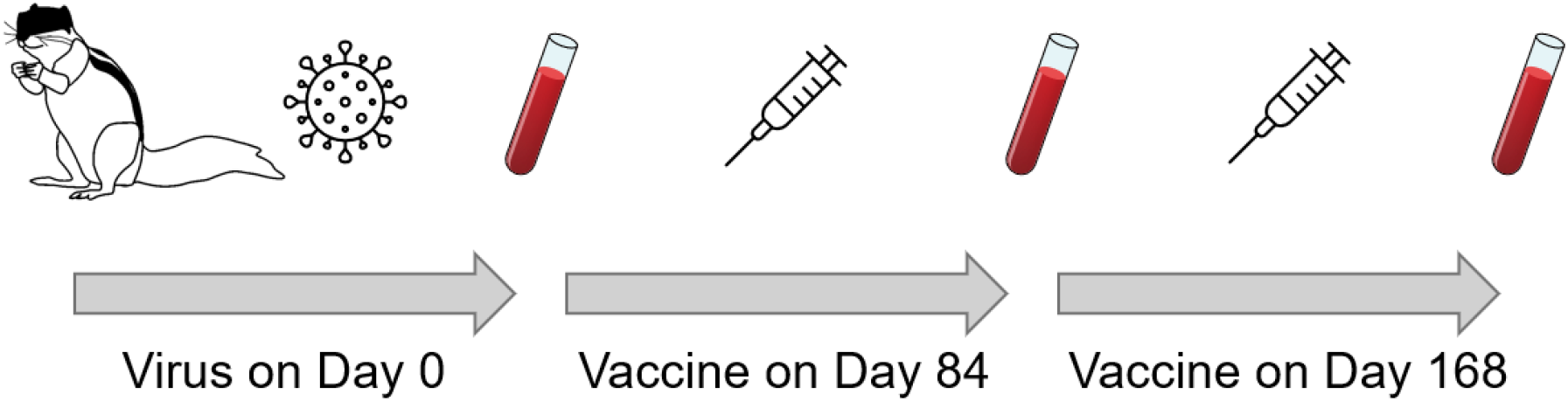
Flowchart of the ferret trial [14]. The influenza virus was inoculated into each animal on Day 0, followed by two vaccinations on Day 84 and Day 168. Blood samples were collected on Day 14, 98, and 182.

**Figure 2:**
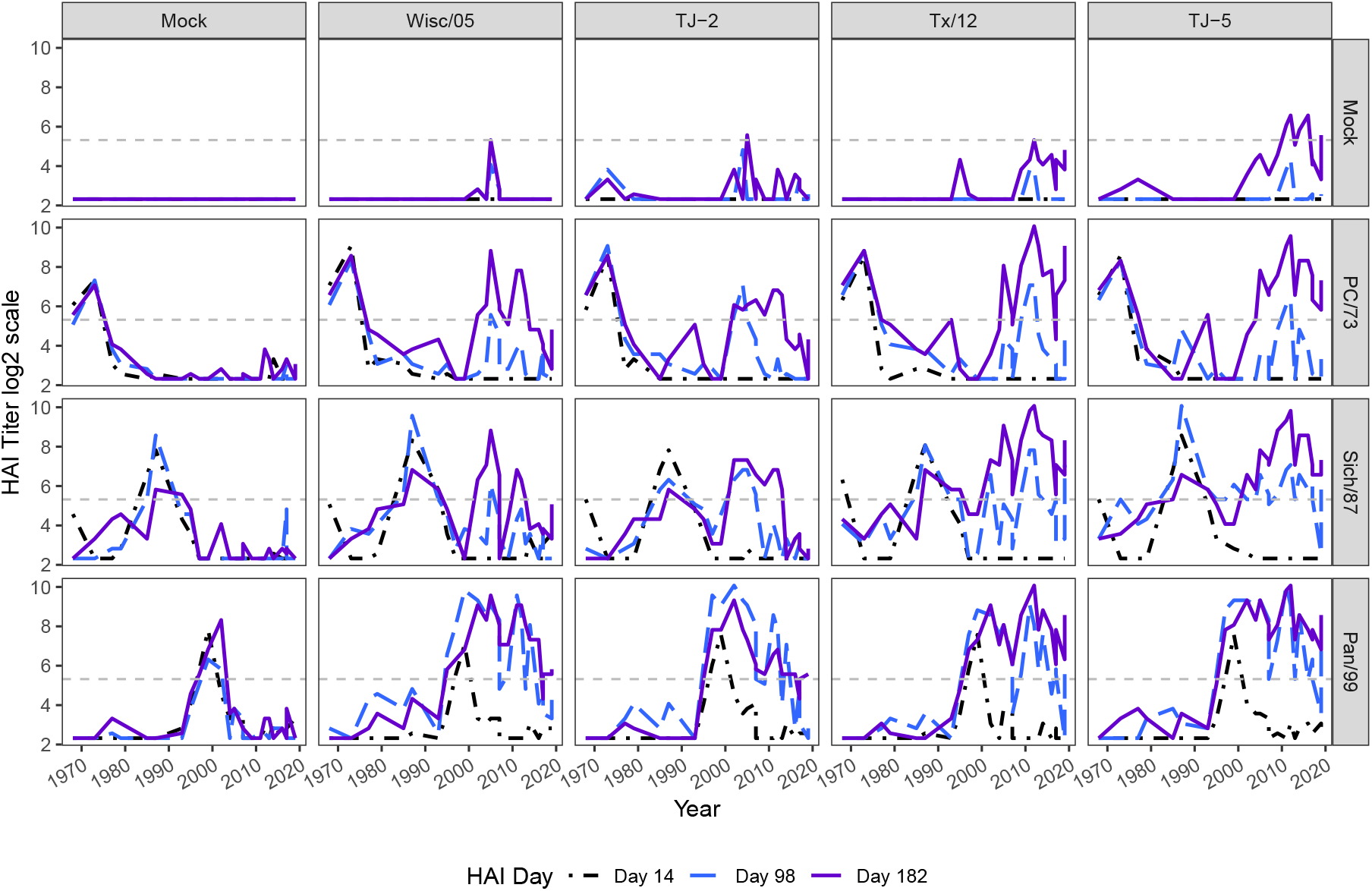
HAI titer of animals against a panel of 25 historical influenza viruses. This figure illustrates the HAI titers of pre-immune and vaccinated animals against 25 historical influenza viruses. A total of 80 ferrets were distributed across 20 groups, each exposed to varying combinations of pre-immunity challenge viruses (right labels) and influenza vaccines (top labels). The x-axis represents the years corresponding to the historical viruses in the HAI assay panel, while the y-axis displays HAI titers on a log2 scale. The data is derived from samples collected at three time points (Day 14, Day 98, and Day 182). Each curve represents the average HAI titers of four ferret samples for each HAI panel strain. The horizontal dash line represents 1:40 in log2 scale (5.32)

**Figure 3:**
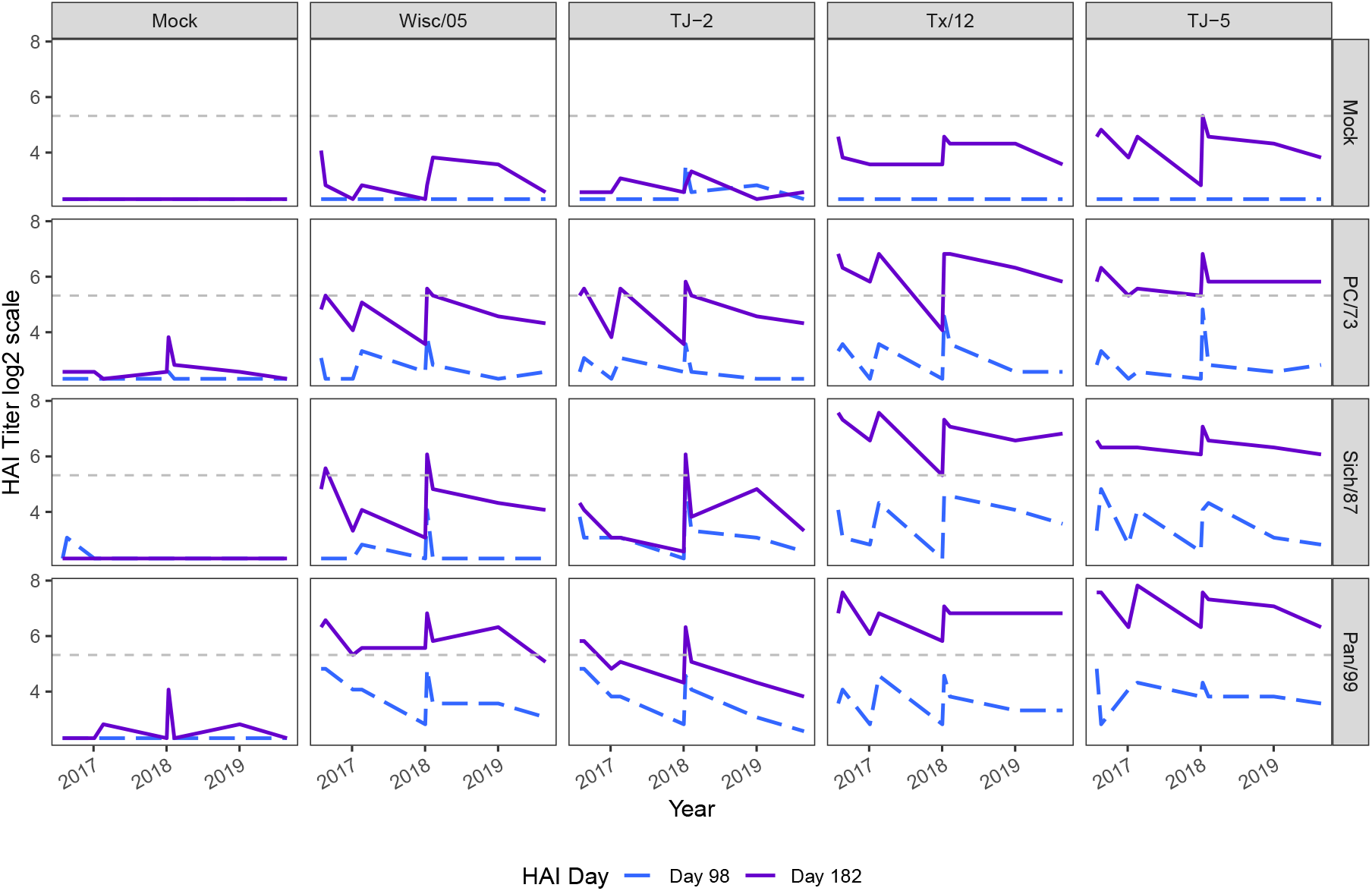
HAI titers of vaccinated animals against 9 co-circulating influenza viruses. This figure illustrates the HAI titers of animals following 1st and 2nd vaccinations against 9 co-circulating influenza viruses. A total of 80 ferrets were distributed across 20 groups, each exposed to varying combinations of pre-immunity challenge viruses and influenza vaccines. The x-axis represents the time corresponding to the co-circulating viruses in the HAI assay panel, while the y-axis displays HAI titers on a *log*_2_ scale. The data was derived from samples collected at two time points (Day 98 and Day 182). Each curve represents the average HAI titers of four ferret samples for each HAI panel strain. The horizontal dash line represents 1:40 in log2 scale (5.32)

In Figure 2, the HAI titer against 25 historical H3N2 strains is presented in chronological order for each of the 20 trial groups. The groups, listed in the first row of the figure, received placebo challenging virus (pre-immunity: mock), while the following three rows represent ferrets infected with one of three influenza viruses. Without pre-immunity, the post-vaccination HAI titers of all the animals on Day 98 were below 1:40 (The top row, Mock: 2.32 in log2 scale, Wisc/05: 2.43, TJ-2: 2.55, Tx/12: 2.44, TJ-5: 2.49). Across all vaccine groups, peaks were observed in years either close to the year that the pre-immunity virus was isolated, or the year the vaccine strain was isolated. For panels that had bigger year differences between the vaccine and pre-immunity, several separate peaks in antibody titer were observed. When the difference was about several years, less peaks were observed.

In Figure 3, similar to the historical figure, the HAI titers of 9 co-circulating strains are presented in chronological order for each of the 20 trial groups. As no data was available on Day 14, we only presented HAI titers on Day 98 (1st vaccine) and Day 182 (2nd vaccine). In general, the pattern was similar to Figure 2. However, the difference in antibody titer between 1st and 2nd vaccination was bigger than the historical strains (Figure 2). The overall mean difference of all pre-immunity groups (log_2_ HAI_*Day*182_ − log_2_ HAI_*Day*98_) were Mock: 0.14, Wisc/05: 1.63, TJ-2: 1.05, Tx/12: 2.73, TJ-5: 2.73. All 1st vaccine induced titers were below 1:40, and 7 out of 20 trials groups (35%) had HAI titers above 1:40 following the second vaccination.

### The impact of pre-existing immunity on HAI titer increase

We assessed the impact of pre-existing immunity by comparing groups having the same vaccine but exposed to different pre-immunity virus. For each trial group, we first tested the overall difference, and then implemented pair-wise comparisons.

Pre-existing immunity influenced the titer increase of all four kinds of vaccines on both homologous and heterologous responses (Figure 4). This figure illustrates the HAI titers of animals following 1st and 2nd vaccinations against 9 co-circulating influenza viruses. The average homologous titer increase of 1st vaccination with Tx/12 was 4.375 (Figure 4, 1st row). For the homologous titer increase (Figure 4, the 1st and 3rd row), all three kinds of pre-existing immunity were associated with a higher 1st vaccine induced titer increase than in the group with no pre-existing immunity. The mean homologous 1st vaccine titer increase was 1.625 in Mock, 4 in PC/73, 4.625 in Sich/87, and 5.75 in Pan/99. However, the 2nd vaccine induced titer increase showed disparate results. The mean homologous 2nd vaccine titer increase was 1.375 in Mock, 3.125 in PC/73, 2.5 in Sich/87, and 1.25 in Pan/99. The homologous titer increase of Tx/12 vaccine were similar across the different pre-existing immunity groups. While, for the Wisc/05 vaccine, the pre-existing immunity of Pan/99 group was associated with slightly lower titer increase compared to the group with no pre-existing immunity (Pan99-Mock = -0.75 titer increase). For the heterologous titer increase (Figure 4, 2nd and 4th row), pre-existing immunity also associated with a higher 1st vaccine induced titer increase as 0.10 in Mock, 0.67 in PC/73, 1.15 in Sich/87, and 2.08 in Pan/99. However, the Pan/99 was associated with slightly lower titer increase in the 2nd Wisc/05, TJ-2, and Tx/12 vaccine.

**Figure 4:**
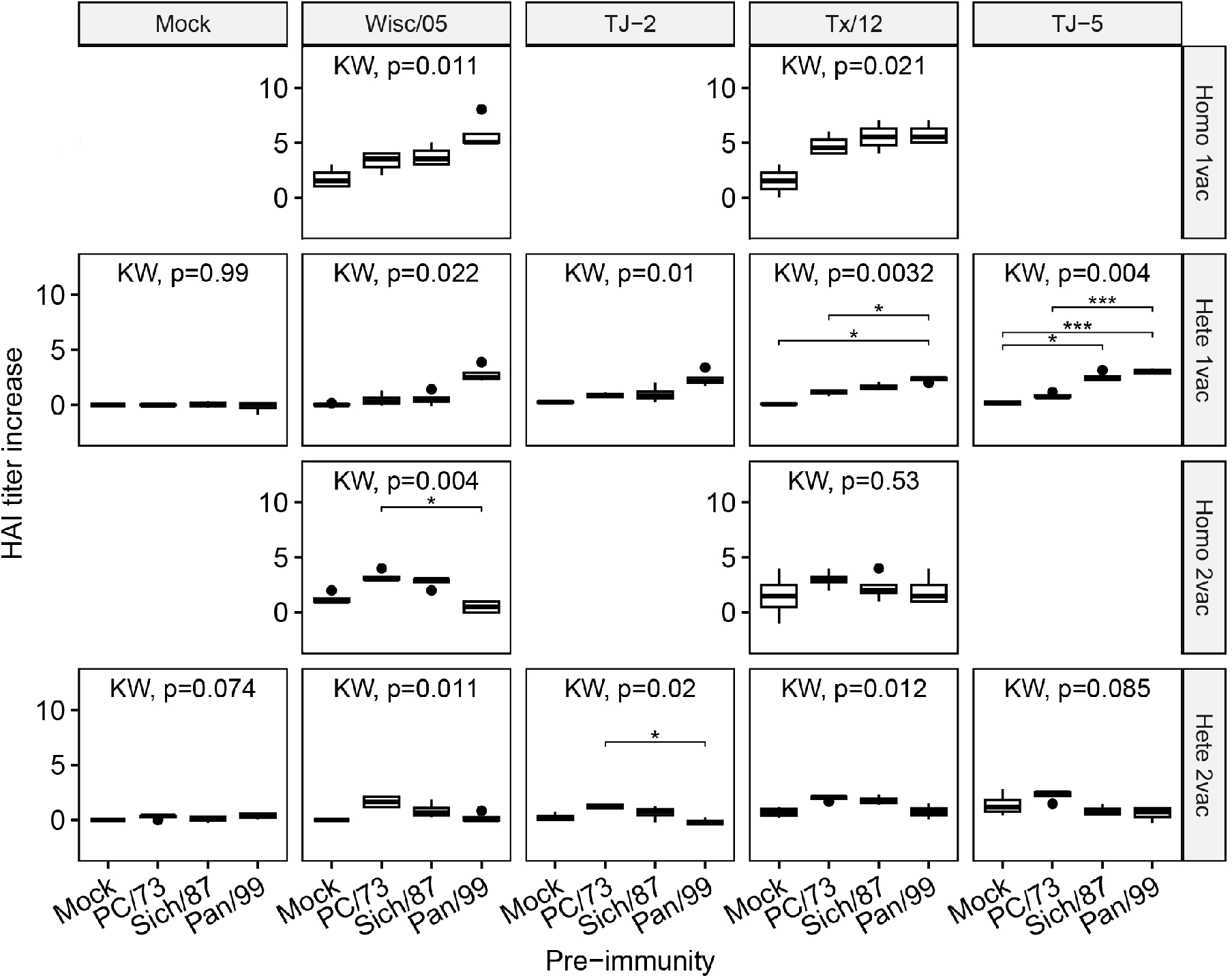
The impact of pre-existing immunity on vaccine induced HAI titer increase (Historical strains). A Kruskal-Wallis (KW) H test was used for comparing if the means of titer increases were same across different pre-existing immunity. We also implemented the Holm-Bonferroni test for pairwise comparisons inside each trial group. For the Holm-Bonferroni tests, we used notations for statistical significance (* indicates *p* < 0.05, ** for *p* < 0.01, and *** when *p* < 0.001). Both homologous (Homo) and heterologous (Hete) responses of the 1st (1vac) and 2nd (2vac) vaccine were presented. As the Mock, TJ-2, and TJ-5 vaccines only had heterologous responses, their homologous panels are empty.

Pre-existing immunity influenced the titer increase of some trial groups on heterologous co-circulating strains (Figure 5). We only had the 2nd vaccine titer increase data of co-circulating strains. The differences in titer increase for TJ-2 vaccine were statistically significant across four kinds of pre-existing immunity settings. However, no significant differences were observed in the other four vaccine groups.

**Figure 5:**
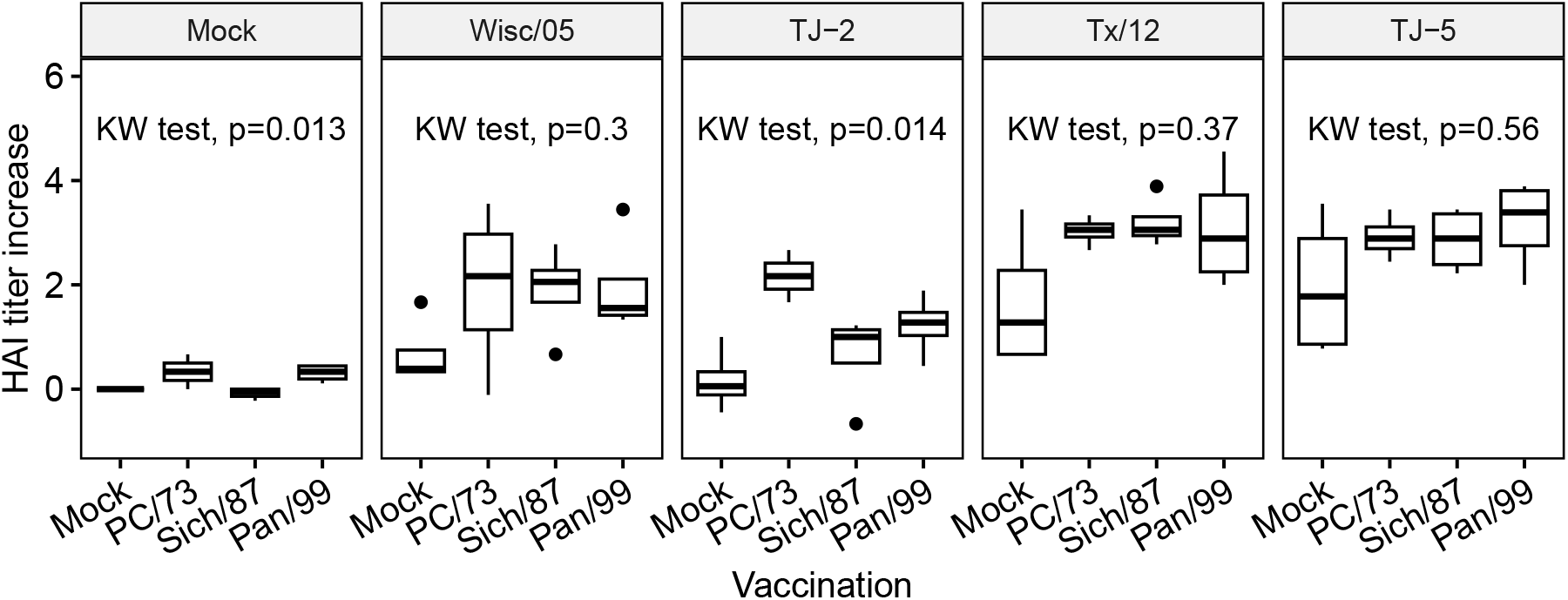
The impact of pre-existing immunity on vaccine induced HAI titer increase (Co-circulating strains). A Kruskal-Wallis (KW) H test was used for comparing if the means of titer increases were same across different pre-existing immunity. We also implemented the Holm-Bonferroni test for pairwise comparisons inside each trial group, but did not find any significant pairs. These co-circulating strains were all heterologous responses.

### The impact of pre-existing immunity on HAI post-vaccination titers

We conducted similar analyses on the HAI titer after the 1st and 2nd vaccinations (Figure 6 and 7). The impacts of pre-existing immunity were statistically significant among most trial groups. Based on pair-wise comparisons, we found the impact of PC/73 and Pan/99 were different for both Tx/12 and TJ-5 vaccine (The 2nd row of the panel). Meanwhile, the mean homologous post-vaccination titers after the 1st vaccination (Wisc/05: 4.07, Tx/12: 3.82) were lower than 5.3 (1:40) when no pre-immunity was given (Mock), but largely improved after the 2nd vaccination (Wisco/05: 5.32, Tx/12: 5.32). However, the heterologous post-vaccination titers were much lower and only slightly increased after the 2nd vaccination (Figure 7).

**Figure 6:**
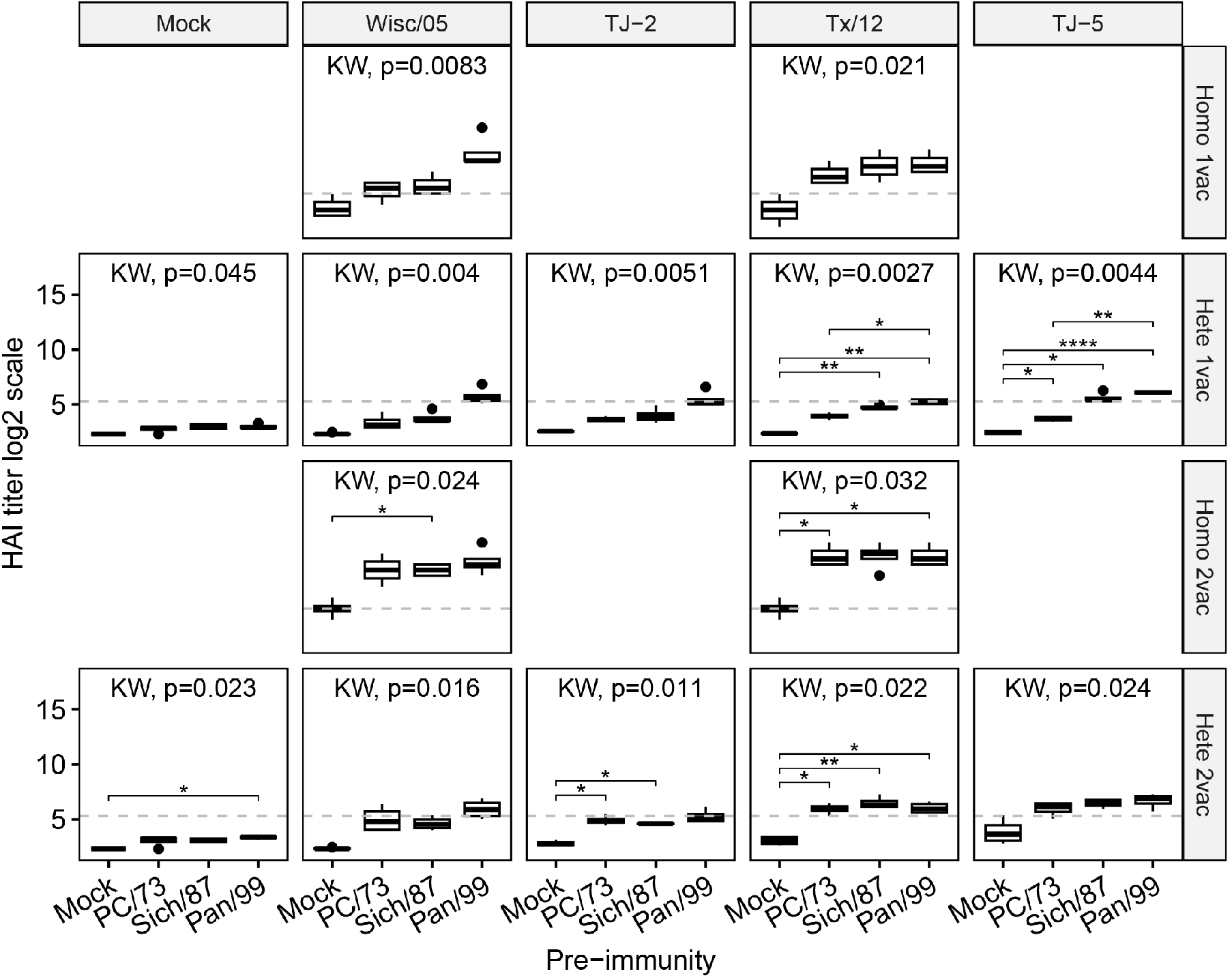
The impact of pre-existing immunity on post-vaccination HAI titer (Historical strains). A Kruskal-Wallis (KW) H test was used for comparing if the means of post-vaccination titer were same across different pre-existing immunity. We also implemented the Holm-Bonferroni test for pairwise comparisons inside each trial group. For the Holm-Bonferroni tests, we used notations for statistical significance (* indicates *p* < 0.05, ** for *p* < 0.01, and *** when *p* < 0.001). Both homologous (Homo) and heterologous (Hete) responses of the 1st (1vac) and 2nd (2vac) vaccine were presented. As the Mock, TJ-2, and TJ-5 vaccines only had heterologous responses, their homologous panels are empty. The horizontal dash line represents 1:40 in log2 scale (5.32)

**Figure 7:**
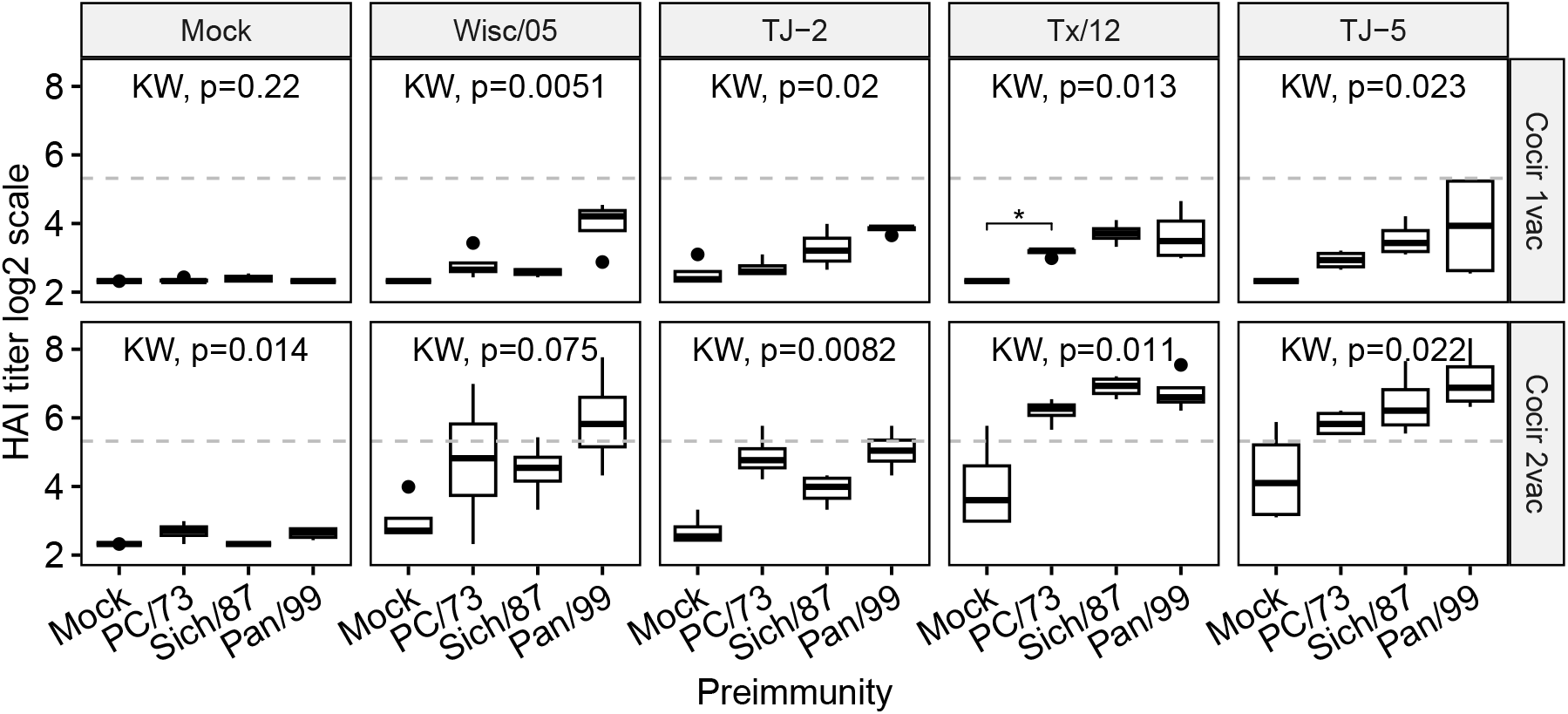
The impact of pre-existing immunity on HAI titer (Co-circulating strains) after 1st (1vac) and 2nd (2vac) vaccine. A Kruskal-Wallis (KW) H test was used for comparing if the post-vaccination titer were same across different pre-existing immunity. We also implemented the Holm-Bonferroni test for pairwise comparisons inside each trial group, but did not find any significant pairs. These co-circulating strains were all heterologous responses. The horizontal dash line represents 1:40 in log2 scale (5.32)

## Discussion

We used HAI data from an animal trial to study the impact of pre-existing immunity on the strength and breadth of the antibody-based immunogenicity of four different vaccines. We observed that different pre-existing immunities led to stronger vaccine induced immune responses, but the magnitude of this change on historical and co-circulating strains varied greatly across groups. In the absence of pre-existing immunity, ferrets generally achieved a comparable level of HAI titer to their counterparts in the pre-existing immunity group with an additional vaccination. Finally, the use of computationally optimized broadly reactive antigens (COBRA), especially TJ-5 vaccines, appears promising in eliciting a broader range of immune responses than traditional wild-type strain vaccination (Appendix).

Pre-existing antibodies due to past infection could interfere with the recognition of immune B cells to new influenza strains, decreasing the response to the latest antigen. However, in Figure 2 and 3, we did not observe this pre-existing immunity related epitope masking effect [22], instead we found that the HAI titer in trial groups having PC/73 or Sich/87 pre-existing immunity had a limited boost of the exposure strain itself. While the vaccine induced responses in trial groups having pre-existing immunity were much higher than naive group (pre-existing immunity equal to mock). Therefore, instead of the theory of “Original Antigenic Sin” which suggesting negative impacts of past exposure on vaccine efficacy [23], we found that pre-existing immunity benefits a vaccine’s elicited protection. There could be an exposure-response association between the pre-infection strain and the later vaccine strain, partially dependent on their similarities. If both strains share a certain number of epitopes, a boost impact can be anticipated. Our results also suggested that the pre-existing antibody levels were maintained at high levels consistent across different vaccines [24].

Pre-existing immunity was associated with better vaccine induced homologous and heterologous immune responses. Compared to the group with no history of pre-immunity exposure, both of our outcomes, HAI titer increase and post-vaccination titer, suggested that pre-existing immunity improved vaccine-induced homologous and heterologous protection (Figure 4 to 7). These differences were more obvious on heterologous protection against historical strains, but less on homologous protection and heterologous protection against modern cocirculating strains. Our results suggested that HA exposures through vaccines among hosts with pre-existing immunity may still be able to create broad protection. We noticed that the theory of “re-set” was reported, suggesting that infections during childhood and adolescence may alter their previous imprinting status [25]. However, we remain unclear about the specific mechanism involved.

Pre-existing immunity is crucial for ferrets in this study to achieve a protective amount of HAI titer with fewer vaccinations. In the trial group with no pre-existing immunity, after the 1st vaccination, we observed that most post-vaccination titer results were below 5.3 = *log*_2_(40), an arbitrary criterion of protection. However, this limitation was almost eliminated after the 2nd vaccination. This result aligns with human vaccination studies suggesting that vaccine-induced protection is limited in toddler or preschooler children [26]. Therefore, considering that many individuals younger than 3 years old have limited pre-existing immunity [27], we support the Recommendations of the Advisory Committee on Immunization Practices, which recommend that certain children aged 6 months through 8 years require 2 doses of influenza vaccine [28].

This animal trial provided valuable insights into understanding the impact of pre-existing immunity on later vaccine induced protections, addressing a challenge in human populations. However, the use of an animal model, ferrets in this study, introduces limitations as the ferret immune system may not be the same as that of humans. Additionally, the relatively small sample size in each trial group may result in low statistical power. Furthermore, the challenge of correlating the age of ferrets to that of humans raises uncertainties, warranting caution in interpreting results for children. Humans also have a more diverse history of infections and vaccinations than our animal model. Therefore, translating our findings to human populations needs further investigations.

In conclusion, our study suggests that pre-existing immunity may strengthen and extend the homologous and heterologous vaccine protection. Individuals without pre-existing immunity may require more vaccinations for optimal protection.

