## Supplemental File for "Pre-existing Immunity to Influenza Aids Ferrets in Developing Stronger and Broader Vaccine-induced Antibody Responses"

### Further data description

Our animal trial data [1] had HAI panel including 25 historical viral strains and 9 co-circulating strains (Table 1 to 2).

Table 1: The name list of 25 historical viral strains

| Historical Strain Full Names | Abbreviations |
|---|---|
| A/Hong Kong/8/1968 | HK/68 |
| A/Port Chalmers/1/1973 | PC/73 |
| A/Texas/1/1977 | TX/77 |
| A/Bangkok/1/1979 | Bgk/79 |
| A/Mississippi/1/1985 | Miss/85 |
| A/Sichuan/2/1987 | Sich/87 |
| A/Shandong/9/1993 | Shan/93 |
| A/Nanchang/933/1995 | Nan/95 |
| A/Sydney/05/1997 | Syd/97 |
| A/Panama/2007/1999 | Pan/99 |
| A/Fujian/411/2002 | Fuj/02 |
| A/New York/55/2004 | NY/2004 |
| A/Wisconsin/67/2005 | Wisc/05 |
| A/Brisbane/10/2007 | Bris/07 |
| A/Uruguay/716/2007 | Uru/07 |
| A/Perth/16/2009 | Perth/09 |
| A/Victoria/361/2011 | Vic/11 |
| A/Texas/50/2012 | Tx/12 |
| A/Switzerland/9715293/2013 | Switz/13 |
| A/Hong Kong/4801/2014 | HK/14 |
| A/Singapore/IFNIMH-16-0019/2016 | Sing/16 |
| A/Kansas/14/2017 | Ks/17 |
| A/Switzerland/8060/2017 | Switz/17 |
| A/South Australia/34/2019 | SA/19 |
| A/Hong Kong/2671/2019 | HK/19 |

Table 2: The name list of 9 co-circulating strains

| Co-circulating Strain Full Names | Abbreviations |
|---|---|
| A/Fiji/110/2016 | Fiji/2016 |
| A/Georgia/12/2018 | Georgia/18 |
| A/Nevada/37/2016 | Nevada/16 |
| A/Abu Dhabi/240/2018 | Abu/18 |
| A/Washington/50/2017 | Wash/17 |
| A/Colombia/0082/2019 | Colombia/19 |
| A/Louisiana/39/2019 | Louisiana/19 |
| A/Sao Paulo/690385/2018 | Sao Paulo/18 |
| A/Moscow/41/2017 | Moscow/17 |

### Additional results

In the main manuscript, we compared one vaccine induced HAI titers across different pre-immunity settings. In the additional results, we compared five vaccine induced HAI titers across the same pre-immunity settings for vaccine comparisons (Figure 1 to 4).

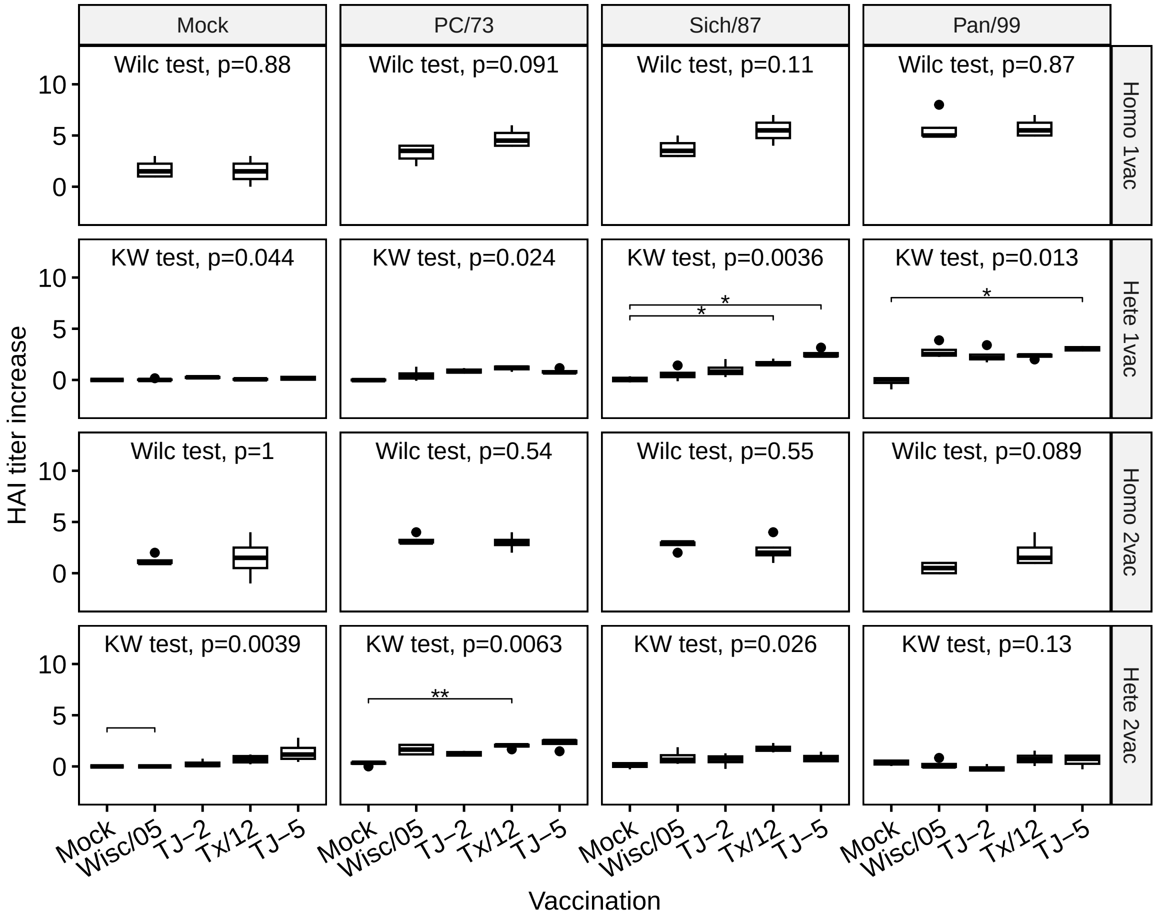


Figure 1: The impact of pre-existing immunity on vaccine induced HAI titer increase (Historical strains). A Kruskal-Wallis (KW) H test was used for comparing if the means of titer increases were the same across different vaccines. We also implemented the Holm-Bonferroni test for pairwise comparisons inside each trial group. For the Holm-Bonferroni tests, we used notations for statistical significance (* indicates $p < 0.05$, ** for $p < 0.01$, and *** when $p < 0.001$). Both homologous (Homo) and heterologous (Hete) responses of the 1st (1vac) and 2nd (2vac) vaccine were presented.The horizontal dash line represents 1:40 in log2 scale (5.32)

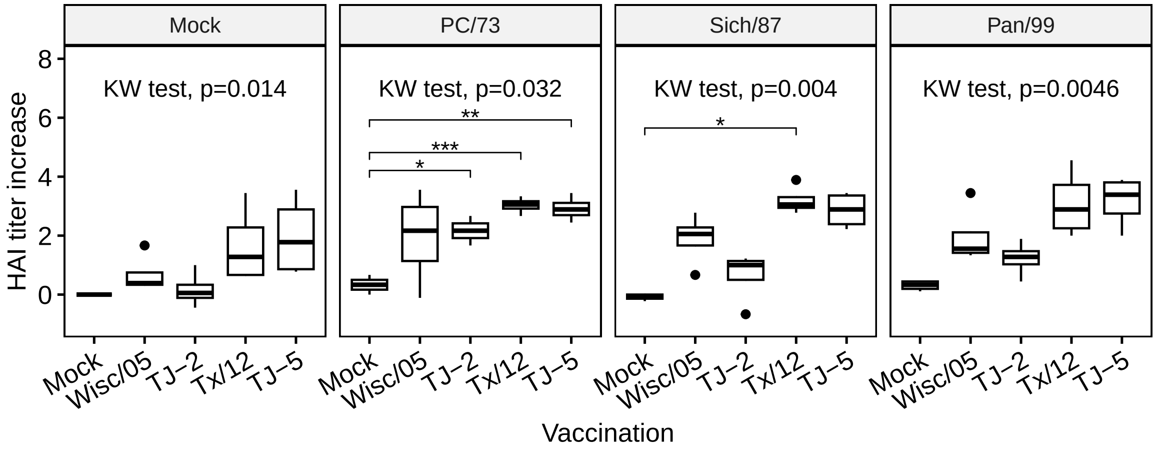


Figure 2: The impact of pre-existing immunity on vaccine induced HAI titer increase (Co-circulating strains). A Kruskal-Wallis (KW) H test was used for comparing if the means of titer increases were the same across different vaccines. We also implemented the Holm-Bonferroni test for pairwise comparisons inside each trial group. For the Holm-Bonferroni tests, we used notations for statistical significance (* indicates $p < 0.05$, ** for $p < 0.01$, and *** when $p < 0.001$). Both homologous (Homo) and heterologous (Hete) responses of the 1st (1vac) and 2nd (2vac) vaccine were presented.The horizontal dash line represents 1:40 in log2 scale (5.32)

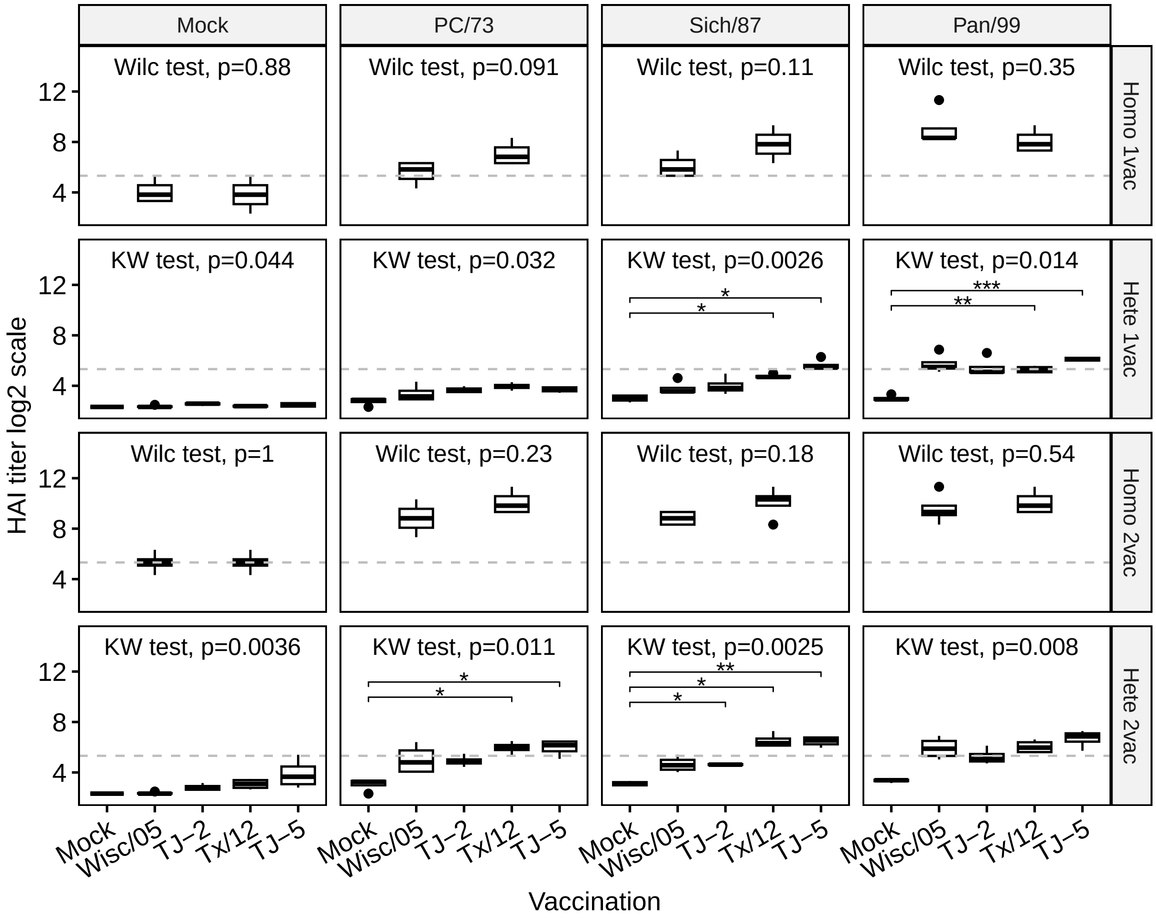


Figure 3: The impact of pre-existing immunity on post-vaccination HAI titer (Historical strains). A Kruskal-Wallis (KW) H test was used for comparing if the means of titer were the same across different vaccines. We also implemented the Holm-Bonferroni test for pairwise comparisons inside each trial group. For the Holm-Bonferroni tests, we used notations for statistical significance (* indicates $p < 0.05$, ** for $p < 0.01$, and *** when $p < 0.001$). Both homologous (Homo) and heterologous (Hete) responses of the 1st (1vac) and 2nd (2vac) vaccine were presented.The horizontal dash line represents 1:40 in log2 scale (5.32)

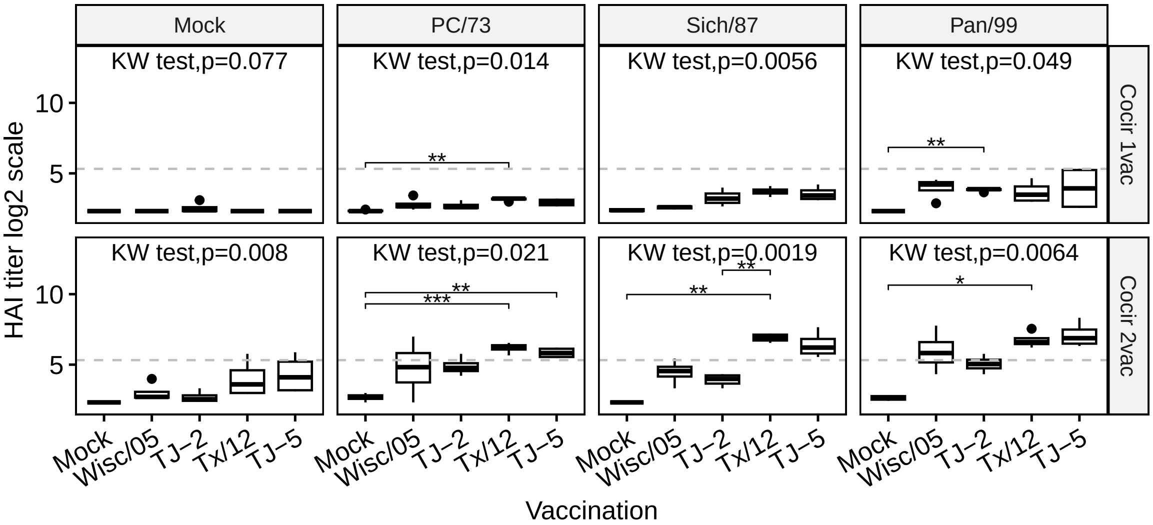


Figure 4: The impact of pre-existing immunity on HAI titer (Co-circulating strains) after 1st (1vac) and 2nd (2vac) vaccine. A Kruskal-Wallis (KW) H test was used for comparing if the means of titer were the same across different vaccines. We also implemented the Holm-Bonferroni test for pairwise comparisons inside each trial group. For the Holm-Bonferroni tests, we used notations for statistical significance (* indicates $p < 0.05$, ** for $p < 0.01$, and *** when $p < 0.001$). Both homologous (Homo) and heterologous (Hete) responses of the 1st (1vac) and 2nd (2vac) vaccine were presented.The horizontal dash line represents 1:40 in log2 scale (5.32)
